# What are you talking about? Representation of the Topic of Speech in the Human Brain

**DOI:** 10.1101/2024.12.12.628085

**Authors:** Adam Boncz, Brigitta Tóth, István Winkler

**Affiliations:** Institute of Cognitive Neuroscience and Psychology, HUN-REN Research Centre for Natural Sciences; Budapest, 1117, Hungary

**Keywords:** Speech, Topic, Functional connectivity, EEG

## Abstract

A sentence outside context provides very restricted information to the listener. Real understanding requires one to be able to place it within the sequence of sentences forming a coherent “story” and just as importantly, within one’s general knowledge of the “topic”. Here we demonstrate the existence of functional brain networks robustly sensitive to the topic of ca. 6-minute-long coherent newspaper articles. Main network hubs are located in parietal and temporal brain regions. These networks changed very slowly during the course of the article, consistent with their involvement in representing the topic rather than the story aspect of the context. Because even the meaning of words can depend on the topic (see e.g., the word “shuttle” in transportation and weaving) the topic-sensitive network must also be intimately linked with the mental lexicon underlying language comprehension.

**Significance Statement:** When we listen to someone talking, our brain continuously integrates the speech stream with our general knowledge associated with the topic. While this is a central process of comprehension, it is still unclear how the brain maintains the relevant representations. Here we demonstrate the existence of a cortical network, which is sensitive to the topic of the speech just heard. This network may form the neural substrate for “topic representation” in the brain. Understanding how such representations are maintained and accessed would let us reveal the computations enabling rapid linguistic integration, a key process in human communication.

## Introduction

When a language researcher flashes a word, such as “five” on a computer screen, English-speaking participants understand it - in a restricted sense. Certainly, one can extract information from this word. However, in real life, language is typically encountered within a coherent context which defines a richer meaning. Compare the word “five” in isolation with the meaning evoked by its use in the sentence “*Five* will do”. The meaning of the word five is now more specific. However, again, if presented in isolation, we understand the sentence only in a restricted sense, i.e., we don’t know five of what is sufficient, what is the purpose, etc. This is because everyday speech is usually ambiguous, its meaning specified by the whole context (*1*).

In terms of a coherent speech segment, context can be subdivided into two classes. One is the immediate context composed of the speech segment itself up to the current point of time - we refer to this type of context as the *story* (also called the narrative, *2-3*). The story is formed by integrating the content of speech over time, chunking memory traces into a chain of events (*4*). The other type of context consists of long-term knowledge evoked by the overall situation and the story so far. It consists of schemas, personal experiences, and general world knowledge (*5–6*). We refer to this latter type of context as the *topic*. Our sample sentence could be part of negotiating the price of a product at the market, discussing the number of workers needed to do a job, etc. “Story” then refers to the sequence of events leading to the statement in the sample sentence, while “topic” covers what we know about bargaining at a market, about the product, the two people negotiating, etc. Here we focus on how the brain treats *topic* by studying the brain networks differentially sensitive to the topic of relatively long coherent speech segments.

Recent work has focused on how the brain builds up a *stor*y over time by linking the temporal course of the narrative of a coherent speech segment to a large-scale network of higher-order brain regions (*2–3, 7*). In contrast to the *story*, the *topic* should change only rather slowly during a coherent speech segment, because the general background does not differ from sentence to sentence. We assume that maintaining a *topic* activates some of the semantic representations assumed to be implemented by a hierarchically distributed network of cortical areas in the brain (*8*). Thus, one can expect that 1) similar large-scale networks are activated throughout a single-topic speech segment across both listeners and time within the segment and 2) different *topics* activate topologically/topographically different networks, consistently within each listener.

These hypotheses were tested on data recorded for a recently published study (*9*) where participants (*N* = 26) listened to four similar speech recordings of descriptive newspaper articles on different topics (a lake, traffic accidents, animal records, and geography; details of all methods can be found in the published paper and the current Supplementary Materials (SI). Other than the difference in the content, recordings were matched in terms of length (ca. 6 mins), speaker delivery, quality, position within the experimental session, task instructions, and affective valence. Thus, the language of the speech and the basic situation were kept constant, so that representations related to the context outside the topic and speech processing in general should not induce differences in the participants’ brain activity while listening to the different speech recordings.

## Results

The participants’ task was to listen to the recordings and to respond to numerals within the speech segments by pressing a button. Further, they were asked questions following each segment probing their memory. After applying standard preprocessing steps to the electroencephalogram (EEG), source activity was reconstructed, averaged in 62 cortical areas, and filtered into the delta to gamma frequency bands.

EEG based functional connectivity (FC) was estimated across all areas in 10-s long, non-overlapping segments (40 epochs/topic see Fig. 1). The selection of epoch length corresponded to the hypothesized slow-changing nature of the topic representations. FC was calculated as the corrected imaginary component of the phase-locking value (ciPLV) (*10*). To reduce the influence of spurious connections, the FC map of each participant and topic was thresholded with a surrogate data-based approach. For illustration, the group average of thresholded FC maps (calculated from all epochs) is depicted in Fig. 2A.

**Figure 1.**
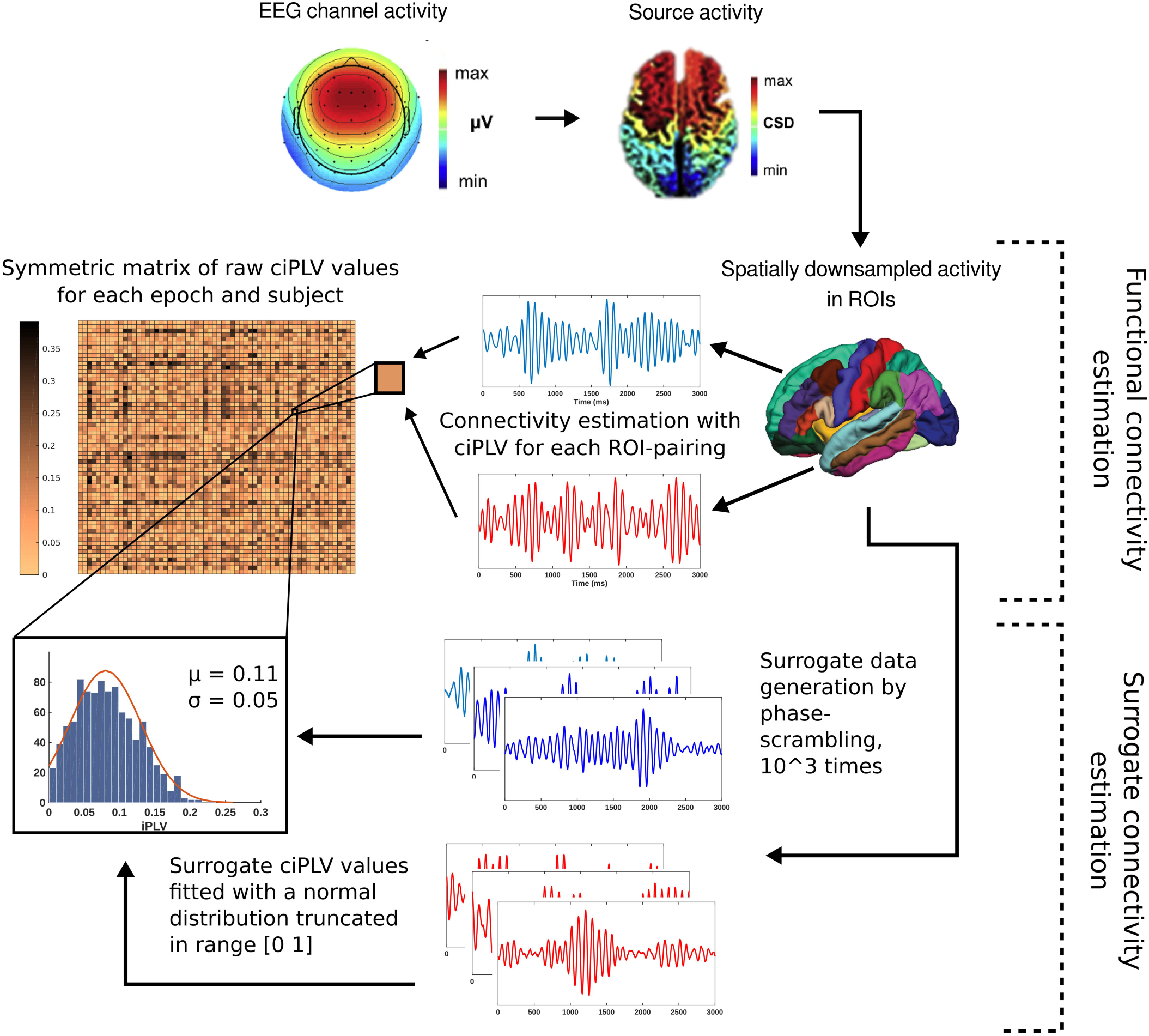
Overview of the FC estimation pipeline. Source-reconstructed signals were first pooled into 62 cortical regions-of-interest (ROIs). Next, for each epoch of each participant, FC for all ROI-pairings was calculated as the corrected imaginary phase locking value (ciPLV), a measure insensitive to volume conduction (zero lag synchronization). For each ROI-pairing, the distribution of surrogate ciPLV values was estimated from 1000 phase-randomized signals. Actual ciPLV values were compared to the surrogate distributions across all epochs with a random permutation test, then corrected with False Discovery Rate (FDR, *q* = .05).

**Figure 2.**
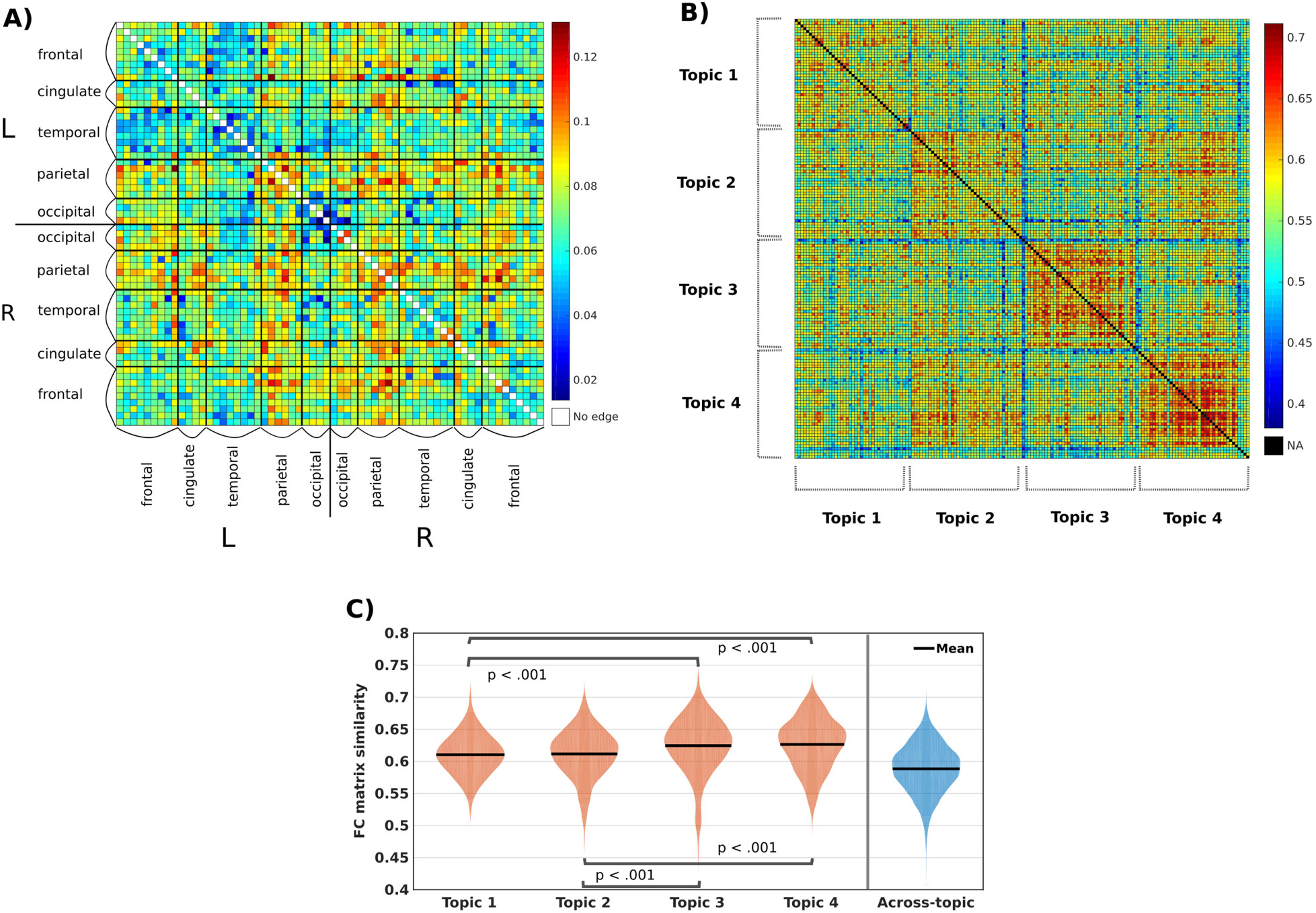
FC matrix similarity within- and across-topics (alpha band). (A) Group average (*N* = 26) FC matrix averaged over all epochs (4 topics × 40 epochs = 160 epochs). Rows and columns represent the 62 ROIs grouped into larger anatomical regions, each cell representing the edge connecting the ROIs of the corresponding column and row. The matrix is symmetric to the diagonal. Color depicts connection strength. Cells with a connection strength not reaching the threshold in any of the participants are marked white. (B) Group average FC matrix similarity values for the four topics. Rows and columns represent epochs, grouped by topic and shown in their order within the speech recording. This matrix is also symmetric to the diagonal. Blue lines at the borders of the topics correspond to the first epoch of the recording. Blocks including the diagonal show the within-topic epoch-pairings. Color depicts similarity strength. (C) Estimated distributions and means of the within-topic FC matrix similarity values, separately for the four topics (pooled from all epoch-pairs of the given topic) and the distribution and means of across-topic FC matrix similarity values (pooled from all epoch-pairs belonging to two different topics). Each within-topic similarity sample differed from the across-topic similarity sample by at least *p* < .001. Bonferroni-corrected significant differences between the four within-topic similarities are marked on the figure.

General topic sensitivity was tested by estimating with Pearson’s correlation the similarity between FC matrices (pairing cells of identical position from the two matrices) across all epochs, both within and across the different speech recordings. This analysis was performed both for the group averages (shown in Figs. 2A and B) and, separately, for each participant. Here we show results for the alpha (8-12 Hz) EEG band, with the other bands, showing similar results, described in SI. The group average FC matrices for epoch-pairings within topics were more similar to each other than for epoch-pairings across topics (random permutation test: *p* < .001; Cohen’s *d* = 0.78). This confirms that FC is affected by the topic of the speech segment. A similar result was obtained for all but three participants’ individual FC matrices (*n* = 23, all *p*s < .004, *d*s > 0.052). Within-topic similarity was also significantly higher than across-topic similarity for each of the four topics (based on the group average FC matrices; *p*s < .001; *d*s > 0.53; see Fig. 2C). Significant similarity differences were also found between the four different topics. Importantly, the FC similarity difference was almost completely independent of performance in the behavioral tasks, except for a small negative relationship between memory performance and similarity difference in the gamma band (see in Table S2).

We also estimated how similarity between group average FC matrices changed in time by regressing the similarity values on the temporal separation between the contrasted epochs. FC similarity was affected by temporal separation (*β* = −0.0007, *t*(3118) = 7.77, *p* < .001). However, topic was a significantly stronger predictor of similarity than temporal separation (likelihood ratio test: χ*²* (1) = 135.08, p < .001 and adjusted R^2^ = 0.019 and R^2^ = 0.06 for models without and with topic as a predictor, respectively). The low rate of change of FC similarity over time (in all frequency bands; see Table S3) supports the notion that the topic consists of a slowly changing set of representations.

For assessing the subset of edges, whose similarity is higher within-than across -topic, the relative contribution of each edge to the difference found for the group-average matrices (Fig. 3A) was evaluated by repeating the similarity analysis for each edge and applying FDR correction (*q* = .05) to control for the large number of statistical tests. *N =* 615 (32.52%) edges showed significant difference for this contrast in the alpha band (for the other bands, see SI). However, this edge selection step is blind to the interactions between edges (i.e., to network properties). Therefore, in order to estimate the smallest network corresponding to topic sensitivity, the edges were ordered according to their effect size (*d*; Fig. 3B) and an Error-Correcting-Output-Codes (*11*) classifier with support vector machine templates was trained on the FC data with the topic label as the output and the number of top contributing edges (*n*) as the step variable. That is, at the *n*th step, the classifier was trained on the *n* individually most contributing edges. Classifier performance at each step was evaluated by the classification error rate (CER) with five-fold cross-validation using 100 repetitions (Fig. 3C). After smoothing mean CER values as a function of edges, a cutoff was applied at the first local minimum, balancing complexity with performance. The final network was then composed of *n* = 29 edges and the final mean CER is 17.33% (Fig. 3D). Based on measures of node centrality (strength, degree, betweenness, and closeness), the most influential nodes of the network are in the left supramarginal, superior parietal, and cuneal areas, as well as in the right inferior parietal gyrus (see SI for the other frequency bands).

**Figure 3.**
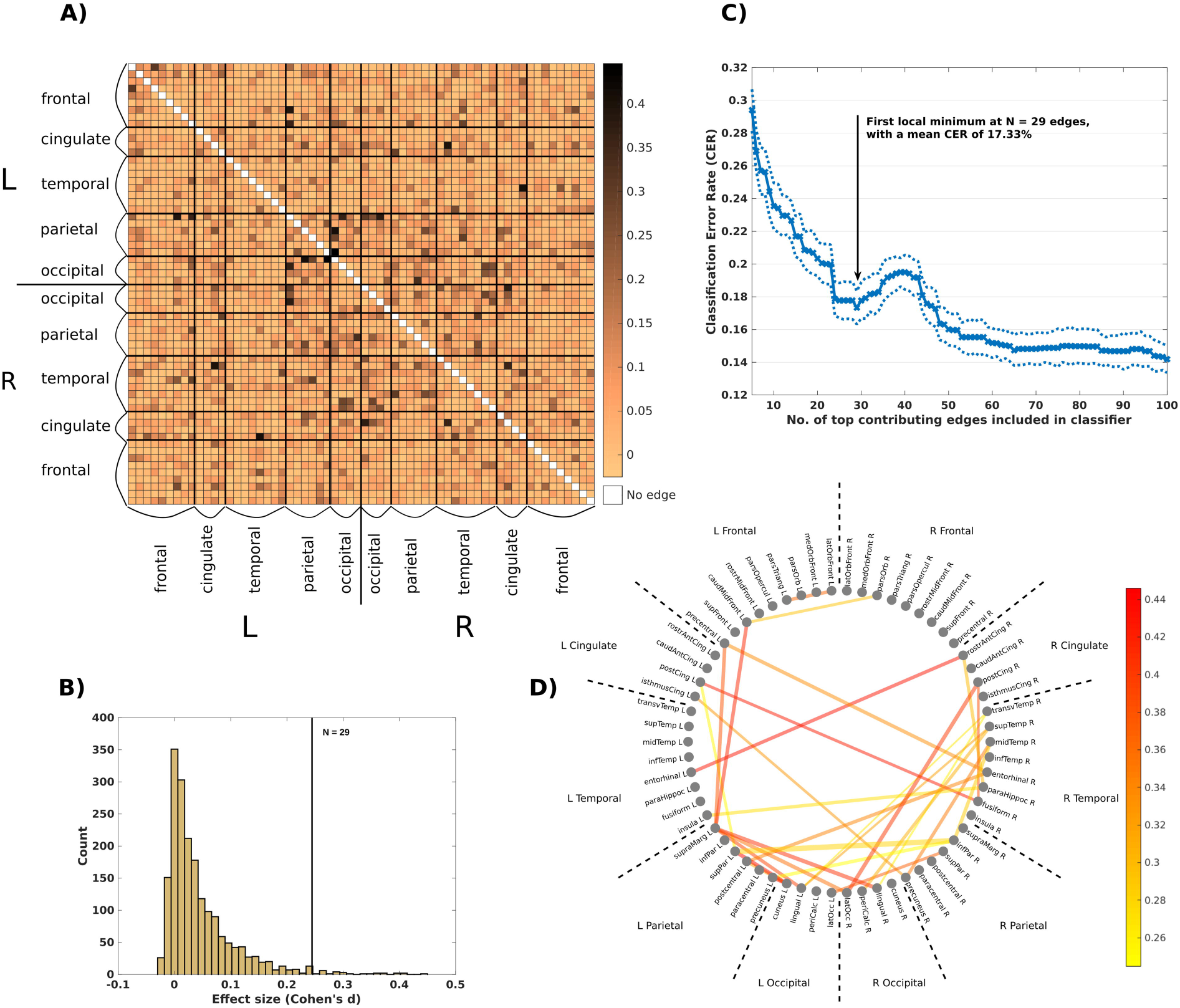
Edge contributions to the FC similarity difference effect (alpha band). (A) Group-averaged (N=26) contribution (effect size, Cohen’s d) of each edge to the group average FC similarity difference between within- and across-topic epoch-pairings, pooled across topics. Rows and columns represent the 62 ROIs grouped into larger anatomical regions. Color scale depicts edge contribution in terms of effect size (d). (B) Histogram of all effect sizes. Vertical line marks the cutoff point (d = 0.247) for the network of edges showing the largest sensitivity (n = 29). (C) Mean CER as a function of the number of edges included in the classifier. The arrow marks the cutoff at n = 29, that is, the first point after which CER increases with more edges added (first local minimum on median filtered CER values). Dotted lines depict the SD of CER values. (D) The 29 edges in the alpha band maximizing sensitivity to the similarity difference between within- and across-topics shown on a circle grouped by anatomical areas. Edge width corresponds to group-level connectivity strength and edge color marks relative contribution to the FC similarity difference between within- and across-topic epochs.

## Discussion

With linguistic and situational variables kept equal, the brain networks extracted from EEG segments recorded while participants listened to spoken newspaper-like articles were significantly more like each other within than across different articles. Robustness of this result is attested by the fact that this relationship was found not only at the group level, but also for ca. 90% of the listeners, individually, as well as separately for each different topic. Similarity was not modulated by the participants’ task performance and the network only changed very slowly over time. These novel results support the assumption that knowledge about the topic is integrated with the linguistic input during speech comprehension and that the representation of the topic is quite stable in time (5, 8).

While the existence of a stable representation of the topic complements the fast build-up of the story, one may ask whether it is separate from the “lexicon”, i.e., the representations assigning meaning to words. If the two are separate, they then must intimately work together towards speech comprehension, as the topic can only be gleaned from an (at least a restricted) meaning of the text, while knowledge of the context is both required to disambiguate certain words and phrases as well as for fuller comprehension. Alternatively, it is possible that context is represented within the lexicon – e.g., word meanings are encoded in relation to their topic (12–13). The latter view is compatible with the notion of a unified memory system (e.g., 14-15) with activation processes preparing items for use. This view receives further support from findings of distributed networks underlying memory-related processes (16–17), including the current study.

The subnetworks sensitive to differences between contexts mainly include parietal and temporal hubs, compatibly with previous descriptions of the topography of semantic and speech processing networks (8, 15). The area with the most consistently central role is the right superior temporal gyrus, ranking among the most central nodes in three separate frequency bands. Note also the right lateralization of central nodes in higher frequencies (beta and gamma). One should, however, consider the topography of the subnetworks with caution, because a) only four specific topics were employed in the current study and b) the topic-sensitive subnetworks somewhat differ across different EEG frequency bands (see SI). The latter result probably reflects that the full network sensitive to the topic includes different types of connections.

In sum, the current data provided strong evidence supporting that a large-scale brain network sensitive to speech topic is activated during speech processing.

## Materials and Methods

Data for the current study was collected as part of an experiment conducted with the aim of studying the effects of the number of speakers on solving the cocktail-party situation (*9*). The full description of the experiment is available in the published paper (*9*) and in the SI, only the condition relevant for the current analyses is detailed here.

### Participants, stimuli and procedures

Twenty-nine native Hungarian speakers (*N* = 29; 11 male; age: *M* = 21.97 years, *SD* = 2.04; 26 right-handed) participated in the study for modest financial compensation. Data from three participants were excluded due to technical errors or extensive motion artefacts (final sample: *N* = 26; 9 male; age: *M* = 22 years, *SD* = 2.15; 24 right-handed).

The speech segments presented to listeners were selected from a larger set of emotionally neutral informative articles collected from news websites. Articles were reviewed by a dramaturg for syntactic pertinence and natural text flow. In the current study, recordings of four articles (on four different topics) were used, voiced by the same male Hungarian actor. The recordings were approximately 6 minutes long.

Speech was delivered to participants at a fixed loudness level of ∼70 dB SPL from a loudspeaker located at 200 cm distance, 30° to the left from midline. The four speech segments were presented in pseudorandomized order, interleaved with other conditions of the full experiment (see in SI).

The experiment was conducted in an acoustically attenuated, electrically shielded, dimly lit room. A 23” monitor was placed at 195 cm in front of the participant, showing an unchanging fixation cross (“+”) during the stimulus blocks. Participants were instructed to avoid eye blinks and other muscle movements and to watch the fixation cross while listening to the speech segments.

Participants performed two tasks concurrently on the speech stream: the “numeral detection” and the “content tracking” task. Details of their procedures, the measured variables, and their relationship with the functional networks of interest are reported in the SI.

### EEG recording and preprocessing

EEG was recorded with a BrainAmp DC 64-channel EEG system with 64 actiCAP active electrodes placed according to the International 10/20 system. The sampling rate was 1 kHz and a low-pass filter with 100 Hz cutoff frequency was applied online. The FCz channel served as the reference electrode during the recording. Bipolar recording from two electrodes placed lateral to the outer canthi of the eyes served for monitoring eye movements. Impedances were kept below 15 kΩ.

EEG data were preprocessed with the EEGlab toolbox (v2019 with Matlab R2017a) (*18*). Offline band-pass filter was applied between 0.5 and 80 Hz by using a finite impulse response (FIR) filter (Kaiser windowed, Kaiser β = 5.65, filter length of 4530) together with another 47.0-53.0 Hz Kaiser bandstop filter (in order to reject noise from the power network). EEG signals were then re-referenced to the average signal of all recorded scalp electrodes. Maximum two malfunctioning EEG channels were interpolated. An automated EOG artifact removal algorithm from the Automatic Artifact Removal toolbox (v1.3) (*19*) was employed, as implemented in EEGlab. The EEG signal was then segmented into 10 s long epochs with 10% overlap between them. The middle 8 s of the first 40 epochs of each trial was used in the subsequent analyses.

### EEG source localization

EEG source reconstruction was performed with Brainstorm (*20*) and followed the protocol of previous studies (*21*, *22*). Forward boundary element head model (BEM) was used as provided by the openMEEG algorithm (*23–25*) and standardized to the MNI brain template (*26*). Default electrode locations were co-registered with the default anatomical mesh. The time-varying source signals (current density) were modeled for all cortical voxels by a minimum norm estimate inverse solution (sLORETA) (*27*). By averaging dipole strengths across voxels, we obtained mean source waveforms (in μAm) for 62 cortical areas based on the anatomical segmentation of the Desikan-Killany-Tourville atlas (*28*). Finally, source activity was filtered into five frequency bands with zero-phase FIR filters (delta: 0.5-4 Hz; theta: 4-8 Hz, alpha: 8-12 Hz, beta: 13-30 Hz, gamma: 30-80 Hz).

For the 64-channel electrode array used here, mean sLORETA localization error has been estimated as 1.40 mm (SD = 3.68) (*22*). Localization was applied identically across participants. Therefore, while overall localization precision might be lower than when individual MRI based anatomical templates are employed, the noise resulting from localization manifests uniformly across the source reconstruction estimates.

### Functional connectivity estimation

All analyses, unless otherwise noted, were carried out with Matlab (R2017a). Details of functional connectivity estimation and subsequent thresholding are depicted in Figure 1.

For each epoch of each participant, functional connectivity (FC) was estimated by calculating the corrected imaginary component of the phase-locking value (ciPLV) (10) from the instantaneous phase of the analytic signals derived from the EEG signals, separately, for each ROI pairings and each frequency band. ciPLV is insensitive to zero-lag couplings arising from signal mixing (*29–30*). FC calculation resulted in one symmetrical (62 × 62) connectivity matrix of raw ciPLV values per participant, epoch, and frequency band.

### Statistical thresholding of functional connectivity values

We tested the connectivity values statistically against surrogate estimates derived from phase-scrambled data (*31-34*). The aim of thresholding was to only retain FC values for the ROI-pairings with larger phase-to-phase coupling strength than that of a linear null model.

First, for each participant, epoch, and frequency-band, we generated 10^3 phase-scrambled versions of the original source signals and calculated ciPLV values across these surrogate source signals. Then, for each of these, we fitted the surrogate ciPLV values with a truncated normal distribution (truncated to the range [0 1], the scale of ciPLV values) and evaluated the fit with the Kolmogorov-Smirnov test. Truncated normal distributions provided good fits to the surrogate data, with rejection rates (at α = .05 level) across all epochs being generally < 0.4% (see in SI). In the subsequent steps, the truncated normal fits were used as surrogate distributions.

The actual connectivity values were compared against the surrogate distributions. That is, for each ROI-pairing, participant, and frequency band, first the surrogate distribution for the average ciPLV value across all 160 epochs was derived from the surrogate distributions calculated for each epoch. Second, the average actual ciPLV value was compared directly against this surrogate distribution, calculating the probability by which the average actual value was larger (stronger FC) than expected from the surrogates. Only the ciPLV values surviving a False Discovery Rate (FDR) correction (*q* = .05) (*35*) were retained. The rest of the connectivity values were set to zero for each corresponding epochs (see the ratios of surviving edges in SI).

The group-average thresholded FC matrix was then calculated for each epoch and frequency band by averaging the thresholded FC matrices across the participants (see Figure 2A for the alpha band).

### Similarity analysis of FC matrix similarities within- vs. across-topics

To test whether group average FC matrices were sensitive to the topic we assessed whether they were more similar across epochs from the same topic than across epochs from different topics. FC matrix similarity across all epoch-pairings was estimated with Pearson correlation. This procedure resulted in a symmetric similarity matrix with a size of 160 × 160 (four topics of 40 epochs, each), separately for each frequency band (see Figure 2B for the alpha band). Within-topic and across-topic similarity values were compared with a random permutation test (with 10^4 permutations) using the studentized (normalized by sample size and variance) mean as test statistic (*36–37*). With studentization, the random permutation test upholds asymptotic validity even in the general case of non-identical distributions underlying the samples (*36*). The random permutation test was then repeated for the four topics separately to test the relationship between within- and across-topic for each individual topic. Finally, to assess whether the relationship found for the group also holds for each individual participant, individual similarity matrices were calculated and compared between within-topic (collapsing the 4 topics) and across-topic (collapsing the 6 different topic pairs).

Within-topic FC matrix similarity values for the group average FC matrices were also compared across the four topics by one-way dependent measures ANOVAs (one for each frequency band) to assess whether the four topics differed from each other in their level of within-topic similarity.

### Testing the relation between temporal separation and FC matrix similarity

We tested the effects of temporal separation (distance) between within-topic epoch-pairings (1-39 for the 40 epochs/topic) on FC matrix similarity of the epoch-pairings with linear models (LM), separately for each frequency band. LMs were fitted with FC matrix similarity as the target variable and either only Temporal Separation as predictor, or both Temporal Separation and Topic as predictors (with no interaction term, as the goal was to assess the effect of Temporal Separation). Temporal Separation was treated as a continuous predictor while Topic as a categorical one. Models were compared with likelihood-ratio tests.

### Edge contributions to FC matrix similarity differences between within- and across-topic

FC matrix similarity was estimated with Pearson correlation which is the sum of edgewise products of the vectorized matrices following z-transformation. For each edge (for a total of 1891 edges in each FC matrix), edgewise products contribute independently to similarity values. Thus, the contribution of each edge to the overall difference between within- and across-topic epoch-pairing similarities can be estimated by repeating the similarity analysis separately for each edge. As for the estimation of the difference between within-topic vs across-topic similarity values, the edge contribution estimation was performed on the thresholded group average FC matrices (one FC matrix for each epoch and topic), separately in each frequency band.

First, each FC matrix was z-transformed so that edgewise products across FC matrices estimate the edge contributions to the correlation (similarity) between the two matrices. Then, for each edge, epoch-pairing similarity values were calculated as the product of the edge values, yielding an epoch-pairings similarity matrix for each edge. Finally, for each edge, using the values in its epoch-pairings similarity matrix, a random permutation test with 10^4 iterations was performed on the difference between within- and across-topic epoch-pairing values with the studentized mean as the test statistic. This procedure resulted in an estimated *p* value and effect size (*d*) for each edge, the latter marking the relative contribution of this edge to the overall similarity difference between the within- and across-topic FC similarity matrices. Due to the large number of comparisons (one for each edge), FDR correction was applied to the significance values (*q* = .05) and only the surviving edges were considered in the classification procedure detailed below.

The topic sensitivity of each edge was tested in isolation in the previous steps. To determine the set of edges, which together maximize the sensitivity of a network, we repeatedly trained and tested the accuracy of a multiclass classifier on the connectivity data of the top 1-300 edges contributing most to the effect in isolation (based on the *d* values calculated in the edge selection procedure). That is, the FC matrices of the epochs were pruned to only contain the connectivity data from the top *n* contributing edges. Then the matrices were vectorized and used as input to the multiclass classifier which was trained to predict the topic label for the connectivity data of each epoch. Error-correcting output codes (ECOC) were used as the multiclass classifier (*11, 38*) (the “fitcecoc” function from the Statistics and Machine Learning Toolbox v11.1 for Matlab). ECOC classifiers use a set of binary classifiers trained in parallel on specific contrasts in the training data. We used linear support vector machine (SVM) templates as the basis of ECOC (46) with one-vs-one coding for a total of six binary classifiers (*39*), one for each of the specific contrasts between two different topics. ECOC performance was evaluated with 5-fold cross-validation. That is, in each run of the classifier, the training data was divided into five sets, and in five rounds, the classifier was trained on four fifth of the data and tested on the remaining part. Overall performance was then determined by the average of the classification error rates (CER) of the five rounds.

At each step of the classification (with connectivity data from each new edge included), the classifier was run 100 times. In order to balance complexity (number of edges included) and performance, the mean CER (average of the 100 runs) as a function of the number of edges included was smoothed with an order 9 median filter and the cutoff was set at the first increase in CER. This results in the first local minimum of CER as a function of the number of edges being selected as the cutoff. The final network comprised of the edges included before the cutoff.

### Node centrality rankings

The relative importance of each ROI in the networks maximizing sensitivity to the within- versus across-topic similarity difference was estimated separately for each frequency band. To this end, the following centrality measures were calculated for each node and frequency band: (1) the number of connections (node degree), (2) the sum of the weights of the connections, (3) betweenness centrality and (4) closeness centrality. These measures were then combined into an aggregate centrality score by z-transforming the values of each centrality measure and averaging them. Nodes with an aggregate centrality score > 1 (meaning at least +1 SD above the mean) are reported as influential (see the results in SI).

## Acknowledgments

The authors are grateful to Gábor Orosz, Ágnes Palotás, and László Hunyadi for text editing, Péter Scherer for voicing the articles, László Liszkai for audio recording and editing, to Zsuzsanna D’Albini and Zsuzsanna Kovács for collecting the EEG data, Zsuzsanna Kocsis for the ICA data preprocessing, Botond Hajdu and Bálint File for providing help in the analysis scripts, and to Eszter Lányi for collecting the study materials for publication.

## Author Contributions

I.W. designed research; A.B. and B.T. performed research and analyzed data; A.B., B.T., and I.W. wrote the paper.

## Data availability statement

All data and code are freely available, and availability is referenced in the Supplementary Materials. Raw EEG has been uploaded to OpenNeuro (openneuro.org/datasets/ds004075). The speech recordings used as stimuli, with their transcriptions, as well as the behavioral data are available at the OSF website of the study at osf.io/56mnz/. Finally, code for connectivity analysis and surrogate data generation is stored at a public GitHubrepo at github.com/dharmatarha/eeg_network_pipeline.

## References

1. U. Hasson, G. Egidi, M. Marelli, R. M. Willems, Grounding the neurobiology of language in first principles: The necessity of non-language-centric explanations for language comprehension. Cognition, 180, 135–157 (2018).

2. C. Baldassano, J. Chen, A. Zadbood, J. W. Pillow, U. Hasson, K. A. Norman, Discovering event structure in continuous narrative perception and memory. Neuron, 95, 709–721 (2017).

3. H. Y. S. Chien, C. J. Honey, Constructing and forgetting temporal context in the human cerebral cortex. Neuron, 106, 675–686 (2020).

4. U. Hasson, J. Chen, C. J. Honey, Hierarchical process memory: memory as an integral component of information processing. Trends Cogn. Sci., 19, 304–313 (2015).

5. P. Hagoort, L. Hald, M. Bastiaansen, K. M. Petersson, Integration of word meaning and world knowledge in language comprehension. Science, 304, 438–441 (2004).

6. K. D. Federmeier, Connecting and considering: Electrophysiology provides insights into comprehension. Psychophysiology, 59, (2022).

7. E. Simony, C. J. Honey, J. Chen, O. Lositsky, Y. Yeshurun, A. Wiesel, U. Hasson, Dynamic reconfiguration of the default mode network during narrative comprehension. Nat. Commun., 7, 1–13 (2016).

8. M. A. L. Ralph, E. Jefferies, K. Patterson, T. T. Rogers, The neural and computational bases of semantic cognition. Nat. Rev. Neurosci., 18, 42–55 (2017).

9. O. Szalárdy et al., https://www.biorxiv.org/content/10.1101/2022.02.21.480990v1 (2022).

10. R. Bruña, F. Maestú, E. Pereda, Phase locking value revisited: teaching new tricks to an old dog. J. Neural Eng., 15, (2018).

11. T. G. Dietterich, G. Bakiri, Solving multiclass learning problems via error-correcting output codes. J. Artif. Intell. Res., 2, 263–286 (1994).

12. A. G. Huth, W. A. De Heer, T. L. Griffiths, F. E. Theunissen, J. L. Gallant, Natural speech reveals the semantic maps that tile human cerebral cortex. Nature, 532, 453–458 (2016).

13. S. Jain, A. Huth, Incorporating context into language encoding models for fMRI. Adv. Neural Inf. Process. Syst., 31 (2018).

14. N. Cowan, Evolving conceptions of memory storage, selective attention, and their mutual constraints within the human information-processing system. Psychol Bull., 104 (1988).

15. J. J. LaRocque, J. A. Lewis-Peacock, B. R. Postle, Multiple neural states of representation in short-term memory? It’s a matter of attention. Front. Hum. Neurosci., 8 (2014).

16. A. J. Watrous, N. Tandon, C. R. Conner, T. Pieters, A. D. Ekstrom, Frequency-specific network connectivity increases underlie accurate spatiotemporal memory retrieval. Nat. Neurosci., 16, 349–356 (2013).

17. S. Hanslmayr, N. Axmacher, C. S. Inman, Modulating human memory via entrainment of brain oscillations. Trends Neurosci., 42, 485–499 (2019).

18. A. Delorme, S. Makeig, EEGLAB: an open source toolbox for analysis of single-trial EEG dynamics including independent component analysis. J. Neurosci. Meth., 134, 9–21 (2014)..

19. G. Gómez-Herrero, Automatic artifact removal (AAR) toolbox v1. 3 (Release 09.12. 2007) for MATLAB (2007), https://germangh.github.io/pubs/aardoc07.pdf

20. F. Tadel, S. Baillet, J. C. Mosher, D. Pantazis, R. M. Leahy, Brainstorm: a user-friendly application for MEG/EEG analysis. Comp. Intell. Neurosci. (2011).

21. B. Tóth, F. Honbolygó, O. Szalárdy, G. Orosz, D. Farkas, I. Winkler, The effects of speech processing units on auditory stream segregation and selective attention in a multi-talker (cocktail party) situation. Cortex, 130, 387–400 (2020).

22. J. Song, C. Davey, C. Poulsen, P. Luu, S. Turovets, E. Anderson, K. Li, D. Tucker, EEG source localization: sensor density and head surface coverage. J. Neurosci. Meth., 256, 9–21 (2015).

23. A. Gramfort, T. Papadopoulo, E. Olivi, M. Clerc, Forward field computation with OpenMEEG. Comp. Intell. Neurosci., (2011).

24. M. Fuchs, R. Drenckhahn, H. Wischmann, M. Wagner, An improved boundary element method for realistic volume-conductor modeling. *IEEE Transact*. Biomed. Eng., 45, 980–997 (1998).

25. M. Fuchs, J. Kastner, M. Wagner, S. Hawes, J. S. Ebersole, A standardized boundary element method volume conductor model. Clin. Neurophys., 113, 702–712 (2002).

26. D. L. Collins, A. P. Zijdenbos, V. Kollokian, J. G. Sled, N. J. Kabani, C. J. Holmes, A. C. Evans, Design and construction of a realistic digital brain phantom. IEEE Transact. Med. Imag., 17, 463–468 (1998).

27. R. D. Pascual-Marqui, Standardized low-resolution brain electromagnetic tomography (sLORETA): technical details. Meth. Find. Exp. Clin. Pharmacol., 24, 5–12 (2002).

28. A. Klein, J. Tourville, 101 labeled brain images and a consistent human cortical labeling protocol. Front. Neurosci., 6 (2012).

29. G. Nolte, O. Bai, L. Wheaton, Z. Mari, S. Vorbach, M. Hallett, Identifying true brain interaction from EEG data using the imaginary part of coherency. Clin. Neurophys., 115, 2292–2307 (2004).

30. J. M. Palva, S. H. Wang, S. Palva, A. Zhigalov, S. Monto, M. J. Brookes, J. M. Schoffelen, K. Jerbi, Ghost interactions in MEG/EEG source space: A note of caution on inter-areal coupling measures. Neuroimage, 173, 632–643 (2018).

31. D. Prichard, J. Theiler, Generating surrogate data for time series with several simultaneously measured variables. Phys. Rev. Lett., 73, 951 (1994).

32. D. S. Bassett, M. A. Porter, N. F. Wymbs, S. T. Grafton, J. M. Carlson, P. J. Mucha, Robust detection of dynamic community structure in networks. Chaos: An Interdisciplinary Journal of Nonlinear Science, 23 (2013).

33. G. C. O’Neill, P. Tewarie, D. Vidaurre, L. Liuzzi, M. W. Woolrich, M. J. Brookes, Dynamics of large-scale electrophysiological networks: A technical review. Neuroimage, 180, 559–576 (2018).

34. A. N. Khambhati, A. E. Sizemore, R. F. Betzel, D. S. Bassett, Modeling and interpreting mesoscale network dynamics. NeuroImage, 180, 337–349 (2018).

35. Y. Benjamini, Y. Hochberg, Controlling the false discovery rate: a practical and powerful approach to multiple testing. J. Roy. Stat. Soc. B Meth., 57, 289–300 (1995).

36. E. Chung, J. P. Romano, Exact and asymptotically robust permutation tests. The Annals of Statistics, 41, 484–507 (2013).

37. A. Janssen, Studentized permutation tests for non-iid hypotheses and the generalized Behrens-Fisher problem. Stat. Prob. Lett., 36, 9–21 (1997).

38. A. Passerini, M. Pontil, P. Frasconi, New results on error correcting output codes of kernel machines. *IEEE Transact*. Neural Netw, 15, 45–54 (2004).

39. J. Fürnkranz, Round robin classification. J. Mach. Learn. Res., 2, 721–747 (2002).

