## Supplementary Information for "What are you talking about? Representation of the Topic of Speech in the Human Brain"

**This PDF file includes:**

Supporting text

Figures S1 to S8

Tables S1 to S4

SI References

Extended methods

As noted in the main text, data for the current study was collected as part of another experiment (*1*). Here we describe the experiment in more detail (including all stimuli and procedures) highlighting the condition analyzed in the current paper. We also provide additional details to the analyses.

**Participants.** Twenty-nine native Hungarian speakers (*N* = 29; 11 male; age: *M* = 21.97 years, *SD* = 2.04; 26 right-handed) participated in the study for modest financial compensation. None of the participants had a history of psychiatric or neurological symptoms. All participants had pure-tone thresholds of <25 dB in the 250 Hz - 4 kHz range, with <10 dB difference between the two ears. Written informed consent was obtained from all participants after the aims and methods of the experiment were explained to them. The study was approved by the United Ethical Committee for Research in Psychology of Hungary (EPKEB), and it was conducted in full compliance with the World Medical Association Helsinki Declaration and all applicable national laws. Data from two participants were excluded from the final analysis due to technical errors, and data from one participant were excluded due to extensive motion artefacts (final sample: N = 26; 9 male; age: *M* = 22 years, *SD* = 2.15; 24 right-handed). Note that the number of participants retained for analysis is higher than that in (*1*), because only a subset of the data was used in the current study.

**Stimuli and procedures.** The speech segments presented to listeners were selected from a larger set of emotionally neutral informative articles collected from news websites. Articles were reviewed by a dramaturg for syntactic pertinence and natural text flow. Twenty-four articles were selected for the recordings. The articles were voiced by two male and one female native Hungarian actor. They were recorded with a Behringer C-1 condenser microphone connected to an Alesis iO2 audio interface sampling the signal at 48 kHz with 32-bit resolution. Recordings were edited offline by a professional audio technician. The average RMS of the sound recordings were equalized to -32dBfs by VST-based attenuation after applying either a -20 dB or -15 dB C3 compressor depending on the dynamics of the recorded speech.

Recording took place in the same room where the experiment was later conducted and the speech segments were played from approximately the same location(s) where the actor(s) sat during the recording session (i.e., the loudspeaker was placed at the approximate location of the actor’s head). Speech was delivered to participants at a fixed loudness level of ~70 dB SPL, using Matlab (R2014a, Mathworks Inc.) with Psychtoolbox 3.0.10 (*2*) running on an Intel Core i5 PC with an ESI Julia 24-bit 192 kHz sound card connected to Mackie MR5 mk3 Powered Studio Monitor loudspeakers.

The experiment was conducted in an acoustically attenuated, electrically shielded, dimly lit room at the Research Centre for Natural Sciences, Budapest, Hungary. Three loudspeakers were placed at an equal 200 cm distance from the participant, positioned symmetrically at -30° (left) 0° (middle), and 30° (right) from midline. Additionally, a 23” monitor was placed at 195 cm in front of the participant, showing an unchanging fixation cross (“+”) during the stimulus blocks. Participants were instructed to avoid eye blinks and other muscle movements and to watch the fixation cross while listening to the speech segments.

In three experimental conditions, one speech stream was delivered alone (Single-speech), or two or three speech streams (One- and Two-distractor, respectively) were presented concurrently. Speech from the left loudspeaker (a male speaker’s speech) was designated as the target of the task (target stream; the only stream in the Single-speech condition). When delivered, the other stream(s) served as the distractor(s). Each condition received four stimulus blocks.

In the current study, only the data collected in the Single-speech condition were analyzed. In this condition, four recordings (on four different topic) voiced by the same male actor were delivered to participants (the same recordings to each participant), in separate stimulus blocks. The audio and the transcripts are publicly available from the Open Science Framework (OSF) website of the study at www.osf.io/56mnz/. The four recordings were approximately 6 minutes long, each (362, 366, 371, and 374 s), delivered with an average speech rate of 271.44 syllables/minute. They included 1719, 1597, 1682, and 1666 syllables and 46, 49, 49, and 50 numeral words (see the participants’ tasks below), respectively.

Participants performed two tasks concurrently on the designated target (in the current context, the only) speech stream: the “numeral detection” and the “content tracking” task. In the numeral detection task, participants were instructed to press a hand-held response key with their right thumb as soon as they detected the presence of a numeral word of 2-4 syllables length. Only numerals indicating the quantity of some object within the context of the text were regarded as targets/distractors. For example, in Hungarian, the indefinite article (“egy”) is the same as the word for “one”. This word, when used as an article, did not constitute a target. There are also words, such as the Hungarian word for daisy (“százszorszép” – literally translated as “hundred-times-beautiful”), which have a numeral as a component. These were not regarded as targets either. These distinctions were explained to listeners prior to the experiment.

For the content tracking task, participants were informed that at the end of each stimulus block, they will be asked to answer 5 questions regarding the contents of the target speech stream. Each question corresponded to a piece of information that appeared within this speech stream. The experimenter read the question and the 4 possible alternative answers. The listener was then asked to verbally indicate his/her choice for the correct answer (multiple-choice test). The experimenter noted the participant’s choice and followed up with a request to the participant to assess his/her confidence in the choice made: “I don’t remember I was just guessing”, “I am not sure, but the option I chose sounded familiar”; I think I heard it during the last block”, “I am sure; I remember having heard it during the last block”, “I know the answer from some other source”. The confidence judgment was recorded by the experimenter.

The stimulus blocks were presented in pseudorandomized order: in the first half of the experimental session (blocks 1-6), each condition was presented two times in random order with the restriction that no condition was immediately repeated; in the second half of the session (blocks 7-12), conditions were presented in reversed order with respect to the first half. Participants were allowed to take a break during the experiment after each stimulus block, and there was a longer mandatory break after the 6th stimulus block. Altogether, the experiment lasted ca. 4 hours.

**Behavioral data analysis.** Analysis of the behavioral data was performed similarly to our previous studies (e.g., *3-4*). For the detection task, button presses for correct responses (hits) were initially collected from a 0-5000 ms interval from the onset of the target event. Responses were then rejected if they were longer than 95% (>1489.3 ms) or shorter than 5% (<432.2 ms) of all initially collected potential target responses (collapsed across all conditions and participants). Note that the limits differ from that in (*1*) due to the different number of participants retained. From the accepted responses, log-normalized reaction times (RT) and hit rates (HR) were calculated for each participant, separately for each stimulus block (topic).

Recognition performance in the content-tracking task was calculated as the percentage of correct responses (PCR), separately for each participant and topic. The sensitivity of the measurement was increased by eliminating items (questions), the response to which was above 95% or below 30% correct overall (pooled across participants and conditions). Responses with a confidence judgment of “I know the answer from some other source” were also dropped from the analysis (only for the given participant). Outside this correction, the confidence judgment data was not considered.

Statistical analysis of participants’ performance across the four topics was performed with separate one-way repeated measures ANOVAs for each measure (within-subject factor: Topic).

**Functional connectivity estimation.** The scripts related to FC estimation and subsequent thresholding are available at https://github.com/dharmatarha/eeg_network_pipeline.

**Statistical thresholding of functional connectivity values.** Truncated normal distributions provided good fits to the surrogate data, with the following rejection rates (at α = .05 level) across all epochs. Delta: *M* = 0.38%, *SD* = 0.15; theta: *M* = 0.12%, *SD* = 0.03; alpha: *M* = 0.23%, *SD* = 0.2; beta: *M* = 0.06%, *SD* = 0.01; gamma: *M* = 0.05%, *SD* = 0.01. Following the comparisons between actual connectivity values and surrogate distributions, after applying FDR, the ratios of surviving edges across the participants were *M* = 86.64%, *SD* = 16.21 for the delta; *M* = 73.31%, *SD* = 23.18 for the theta; *M* = 63.06%, *SD* = 23.00 for the alpha; *M* = 78.26%, *SD* = 15.26 for the beta; and *M* = 73.44%, *SD* = 24.48 for the gamma band.

**Testing the relation between behavioral performance and FC matrix similarity.** We tested whether the FC matrix similarity difference between within- and across-topic values was related to behavioral measures. To this end, we tested by Linear Mixed Models (LMM) the relationship between the effect size of the difference between the within- versus across-topic FC similarity and (separately) the RT, HR, and recognition performance (PCR) values measured for the topic. The effect size was calculated separately for each participant/topic/frequency band serving as the response variable of LMM, with the behavioral measures as predictors, and participant as the random effect. Separate LMMs were fitted for each of the five frequency bands. Regarding the random-effects structure of the models, we fitted LMMs both with and without random slopes for participants (cf. *5*). We report the results for the models yielding the better fit as measured by the Akaike Information Criterion (AIC) and the Bayesian Information Criterion, (BIC) (*6*). Models were fitted with a restricted maximum likelihood approach using the “fitlme” function of the Statistics and Machine Learning Toolbox for Matlab (v11.1).

**Edge contributions to FC matrix similarity differences between within- and across-topic.** In general, after the first step of edge selection described in the main text, a surprisingly large number of edges showed a significant contribution to the difference between within- versus across-topic similarities (e.g. N = 615 for alpha). To verify our approach to the selection of edges, we also conducted a resampling-based thresholding procedure across all edges. The idea was to estimate the standard deviation for the contribution of each edge with a jackknifing resampling method and reject edges where the contribution effect size was not significantly different from zero. First, the standard deviations of the edge contribution effect sizes were estimated with a delete-one jackknife procedure. That is, edge contributions were recalculated for leave-one-out group average FC matrices, once for each participant, yielding one jackknifed edge contribution effect size for each edge. Then, for each edge, a one-sided z-test was performed to test whether the mean effect size (the d value from the randomization test in the previous paragraph) was larger than zero (assuming normality of jackknifed means based on the central limit theorem). Finally, an FDR (*q* = .05) was applied to the significance values and only the edges surviving the FDR were considered to have passed the resampling-based thresholding. Due to computational load, this calculation was only performed for the alpha band. In the alpha band, all edges with a significant contribution found with the first procedure survived this verification step as well. For the other bands, we assumed that this procedure would not substantially modify the set of edges selected in the previous step.

Extended results

**Behavioral results.** A significant Topic effect was found on the RT of the detection task (*F*(3, 75) = 6.99, *p* < .001, *η*^2^ = .073). Pairwise comparisons (Figure S1A) with Bonferroni correction showed significantly longer RTs for topic no. 3 (*M* = 856.5.8 ms, *SD* = 115.6) than no. 1 (*M* = 782.4 ms, *SD* = 92.8; *t*(25) = 4.87, *p* < .001, *d* = 0.95) and no. 2 (*M* = 783.8 ms, *SD* = 120.6; *t*(25) = 3.50, *p* = .01, d = 0.69). There was a similar effect on the HR of the detection task (*F*(3, 75) = 12.95, *p* < .001, *η*^2^ = .222). Pairwise comparisons (Figure S1B) with Bonferroni correction revealed that the HR for topic no. 3 (*M* = 91.21%, *SD* = 4.43%) was significantly lower than for no. 1 (*M* = 97.80%, *SD* = 3.00%; *t*(25) = 7.90, *p* < .001, Cohen’s *d* = 1.55) and no. 2 (M = 95.90%, SD = 4.45%; t(25) = 3.98, *p* < .001, d = 0.78). Further, topic no. 1 yielded a significantly higher HR than no. 4 (*M* = 93.85%, *SD* = 6.25%; *t*(25) = 2.36, *p* = .018, *d* = 0.46). Participants’ recognition memory (*M* = 69.21%, *SD* = 26.06) was significantly above chance level (25% in the 4-alternative forced choice task) for all topics: (all *t*s(25) > 6.06, *p*s < .001, *d*s > 1.19; Figure S1C). No significant effect of Topic was found on recognition memory (*F*(3, 75) = 2.09, *p* = .109, *η*^2^= .054).

**Across- and within-topic FC matrix similarities.** The results for the alpha band are reported in detail in the main text. Here we summarize the results for the rest of the bands (delta, theta, beta, and gamma). There was a significant difference between within- and across-topic FC similarity matrices in all four frequency bands (random permutation tests, all *p*s < .001; see Table S1). The mean connectivity matrices, the FC similarity matrices, and the pairwise comparisons of FC matrix similarity values across topics are depicted in Figures S2-S4.

**Relationship between behavioral measures and FC similarity.** Table S2 contains the results from LMMs testing the relationship between the within- vs. across-topic FC similarity effect and the behavioral measures. Results are reported separately for each frequency band and for each behavioral measure as predictor. The only significant result was in gamma band, with PCR as the predictor (*β* = -0.539, *p* = 0.0105, Bonferroni corrected), suggesting that the similarity effect was smaller in subjects with better memory performance, but only in the gamma band.

**Relationship between temporal separation and FC matrix similarity.** Table S3 depicts the results of LMs testing the relationship between FC matrix similarity and Temporal Separation / Topic. In all frequency bands, we found a significant effect of Temporal Separation. Adding Topic as a factor increased model fit significantly in all cases.

**Topography of the network with higher within- than across-topic similarity.** Following edge selection based on effect sizes and statistical thresholding, the number and ratios of surviving edges for the frequency bands were *N* = 133 (7.03%) for delta; *N* = 69 (3.65%) for theta; *N* = 198 (10.47%) for beta; and *N* = 725 (38.34%) for the gamma band. The final networks selected for maximizing topic sensitivity in the four frequency bands consisted of 65 (delta), 54 (theta), 45 (beta) and 34 (gamma) edges, with mean CER values of 30.53% (delta), 22.76% (theta), 22.08% (beta) and 11.39% (gamma). The edge contributions, CER functions of the number of edges, effect-size histograms, and circular graphs of the selected networks for the four frequency bands are depicted on Figures S5-S8. The most central nodes in each frequency band are reported in Table S4. The full node list with the aggregate centrality values can be found at the OSF website of the study (www.osf.io/56mnz/).

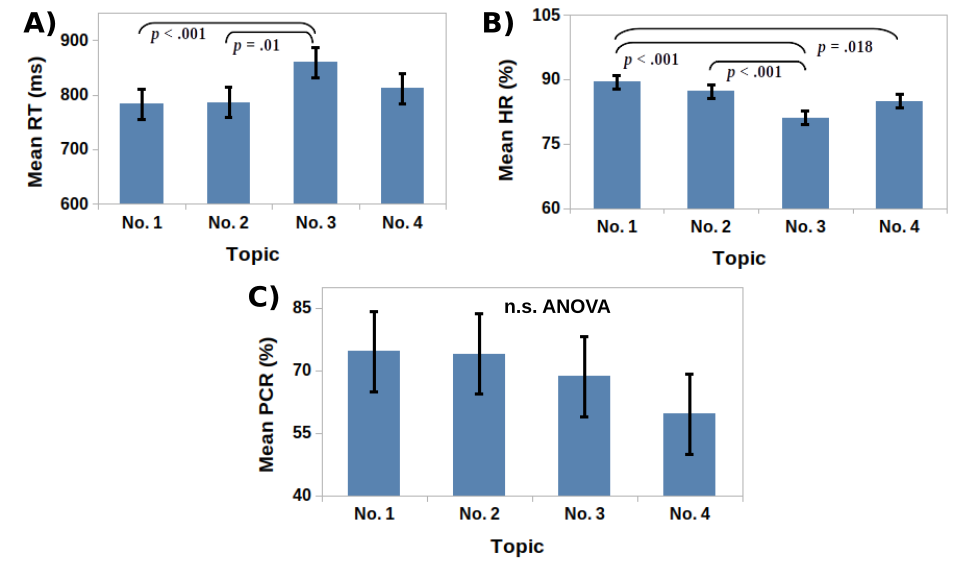
**Fig. S1. Group-averaged (*N* = 26) behavioral measures across the four different topics.** Error bars depict 95% within-subject confidence intervals (*7*). Significant pairwise comparisons (following up significant Topic ANOVA effects) are shown on top of each panel. (**A**) Hit rates (HR); (**B**) Reaction times (RT); (**C**) Recognition memory in percent of the correct responses (PCR).

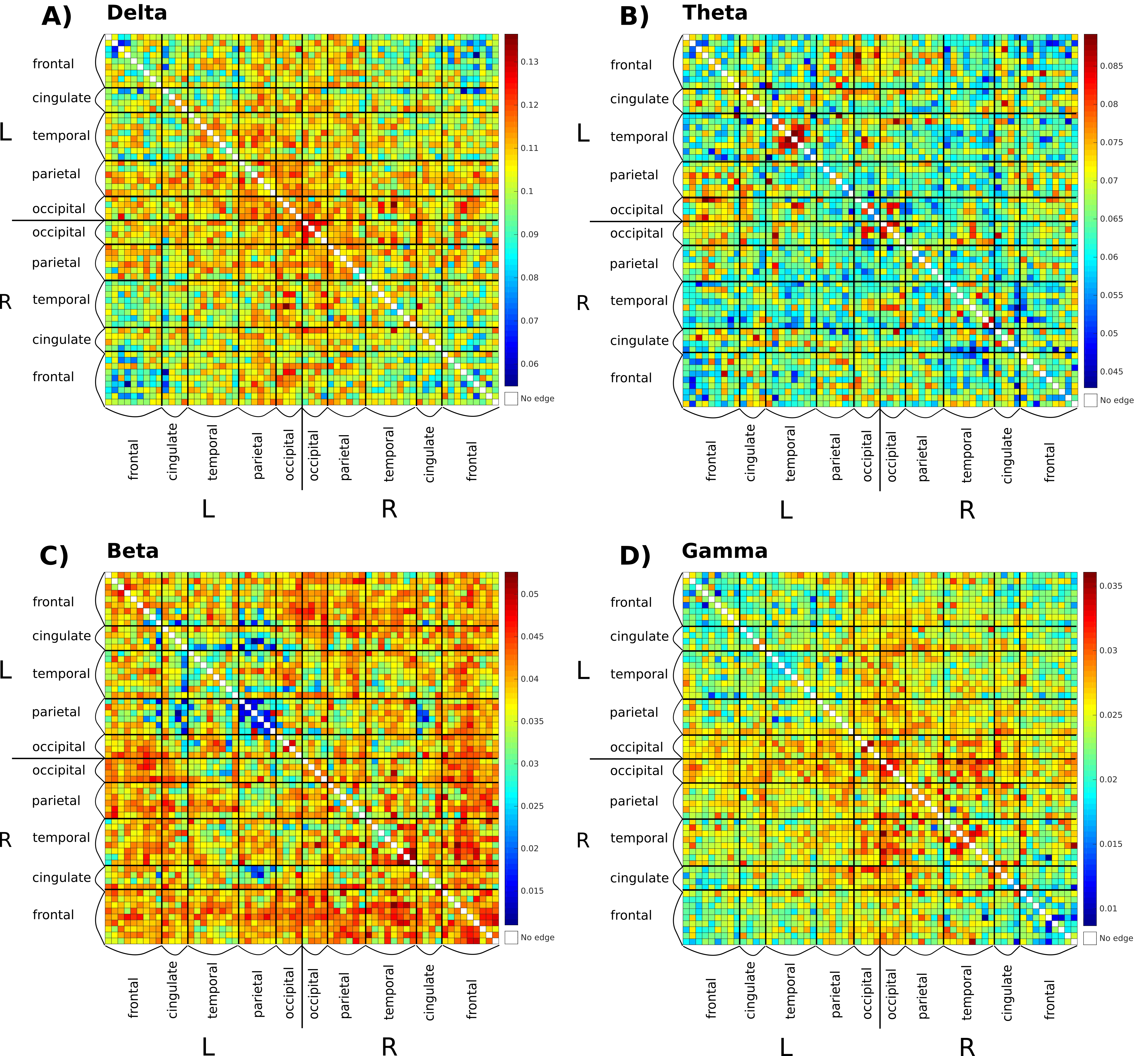
Fig. S2. Group average (*N*=26) FC matrix in the four frequency bands, averaged over all epochs (4 topics × 40 epochs = 160 epochs). Rows and columns represent the 62 ROIs grouped into larger anatomical regions (marked left and below the matrix), thus cells represent an edge connecting the ROI of the column and that of the row. The matrix is symmetric to the diagonal by definition. Color scale for connection strength is at the right side of each panel. Panels: (A) delta; (B) theta; (C) beta; (D) gamma band.

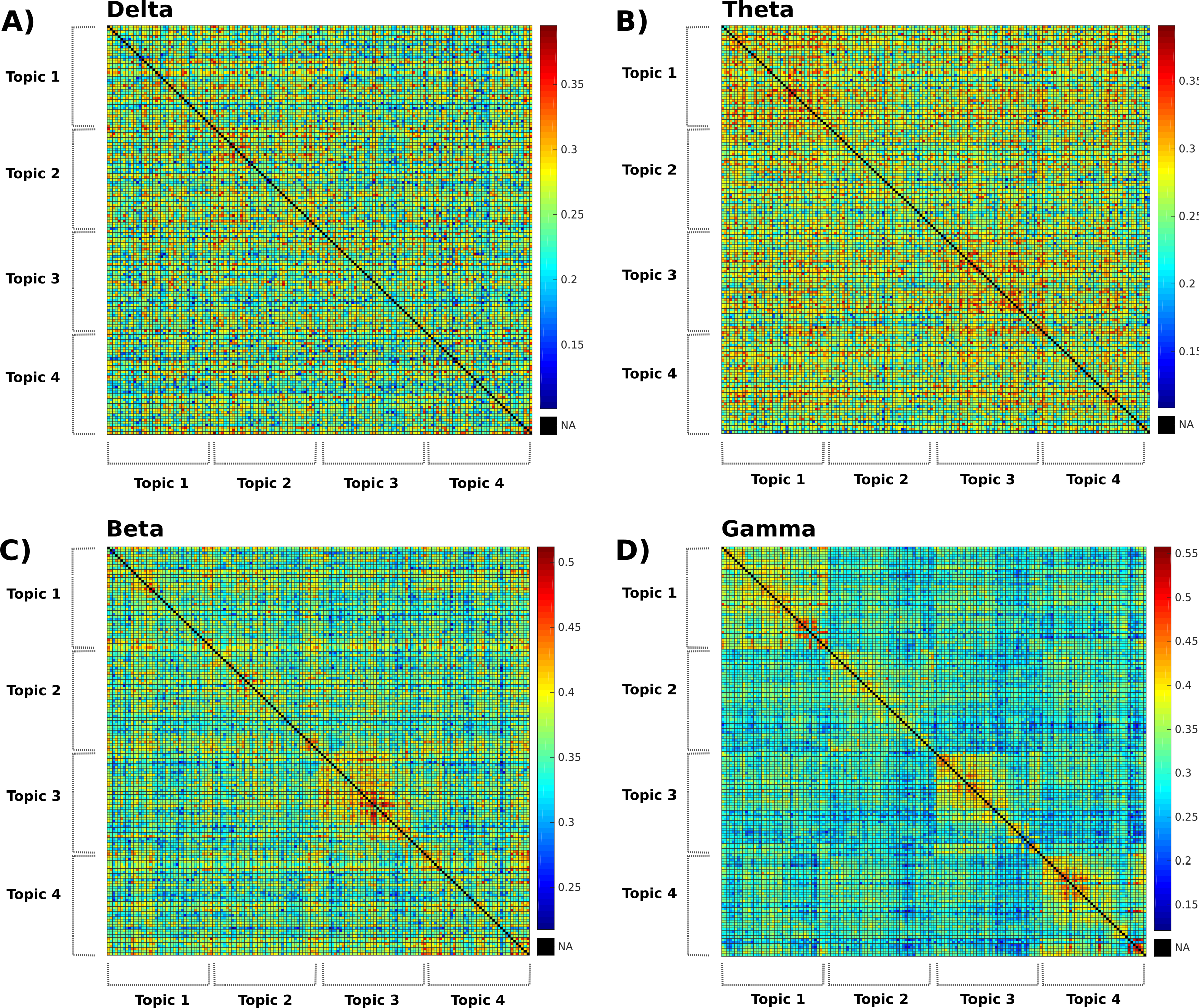

Fig. S3. Group average FC similarity matrices between the four topics, separately for the four frequency bands. Rows and columns represent epochs, grouped by topic (marked left and below the matrix) and shown in the order they appeared within the speech recordings. Panels: (A) delta; (B) theta; (C) beta; (D) gamma band.

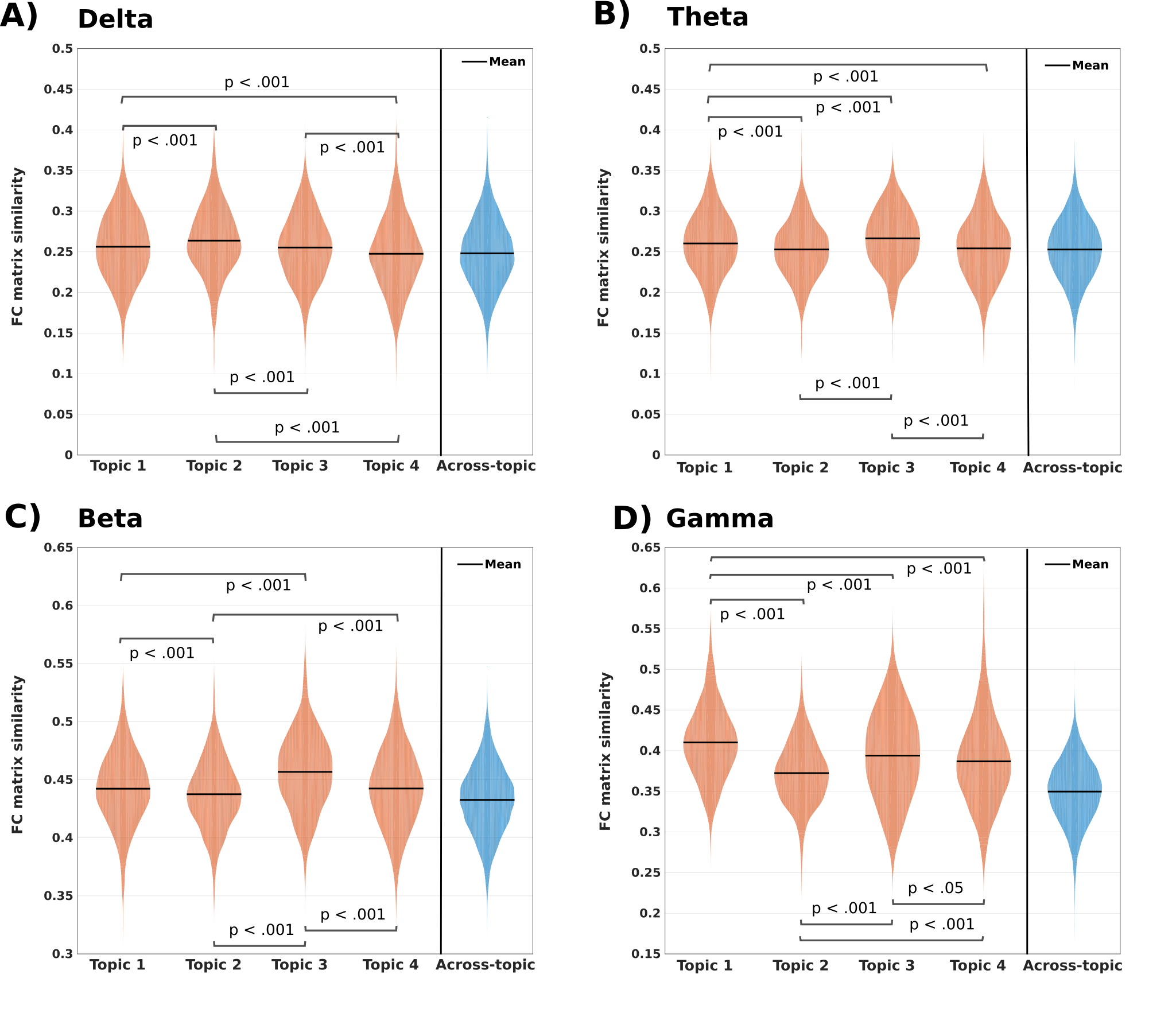

**Fig. S4. Estimated distributions and means of within-topic FC similarity values for the four narratives (pooled from all epoch-pairs of the given topic) and the distribution and mean of across-topic FC similarity values (pooled from all epoch-pairs belonging to two different topics) in the four frequency bands.** Significant differences (Bonferroni corrected within each frequency band) between the within-topic similarities are marked on each figure. Panels: (**A**) delta; (**B**) theta; (**C**) beta; (**D**) gamma band.

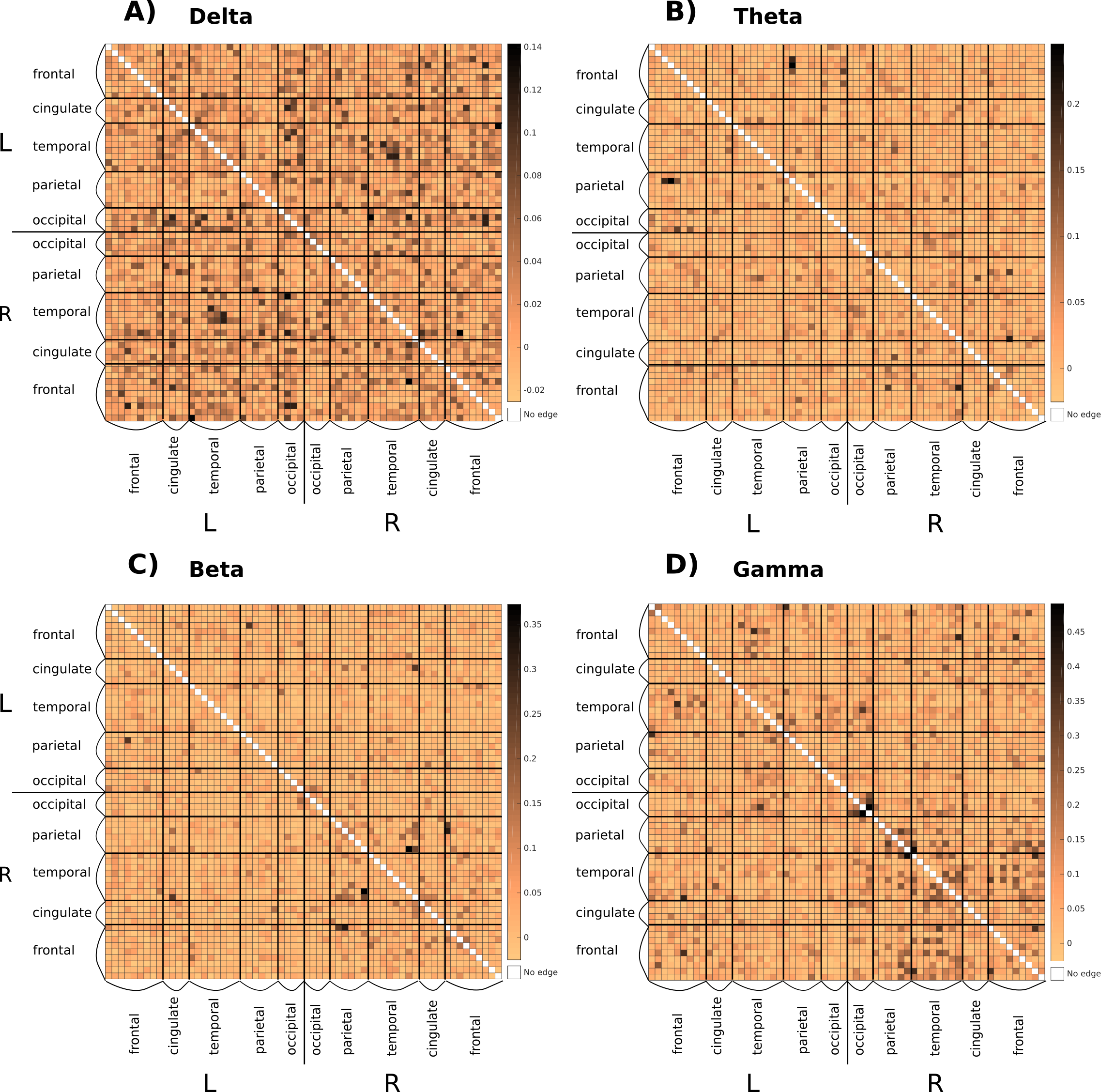
**Fig. S5. Group-averaged (*N*=26) contribution (effect size: Cohen’s *d*) of each edge to the group average FC similarity difference between within- and across-topic epoch-pairings, pooled across topics, depicted separately for the four frequency bands.** Rows and columns represent the 62 ROIs grouped into larger anatomical regions (marked left and below the matrix). Note that color mapping (shown right to each panel) differs across the four frequency bands. Panels: (**A**) delta; (**B**) theta; (**C**) beta; (**D**) gamma band.

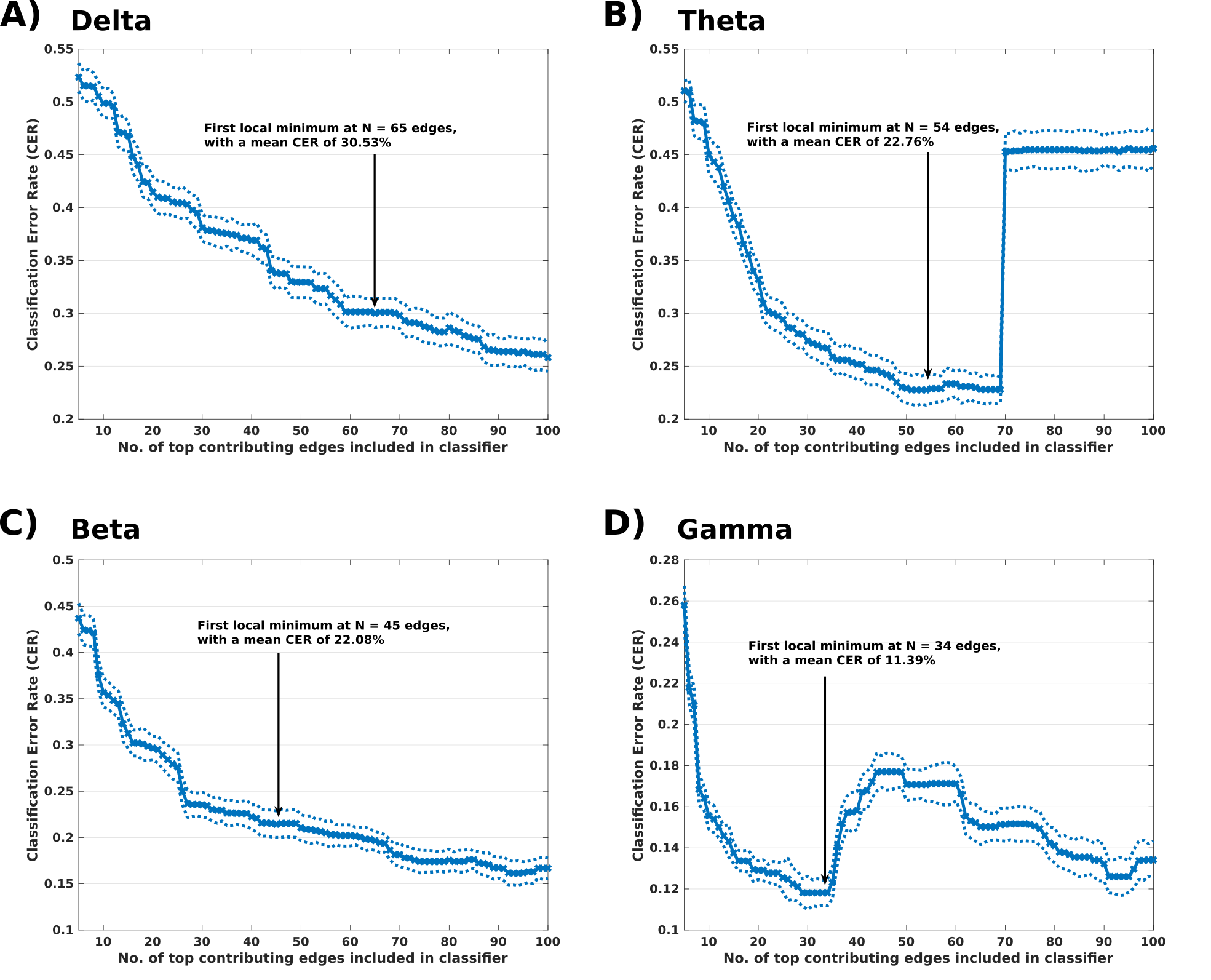

**Fig S6. Classification error rate (CER) as a function of the number of edges included in the classifier for the four frequency bands.** Black arrow marks the cutoff, that is, the first local minimum of the median filtered CER values as a function of the number of edges number. Dotted lines depict the +/- 1 SD range across classifier runs. There were 100 runs for each edge number and frequency band. (**A**) Delta band cutoff is at 65 edges with a mean CER of 30.53%. (**B**) Theta band cutoff is at 54 edges with a mean CER of 22.76%. (**C**) Beta band cutoff is at 45 edges with a mean CER of 22.08%. (**D**) Gamma band cutoff is at 34 edges with a mean CER of 11.39%.

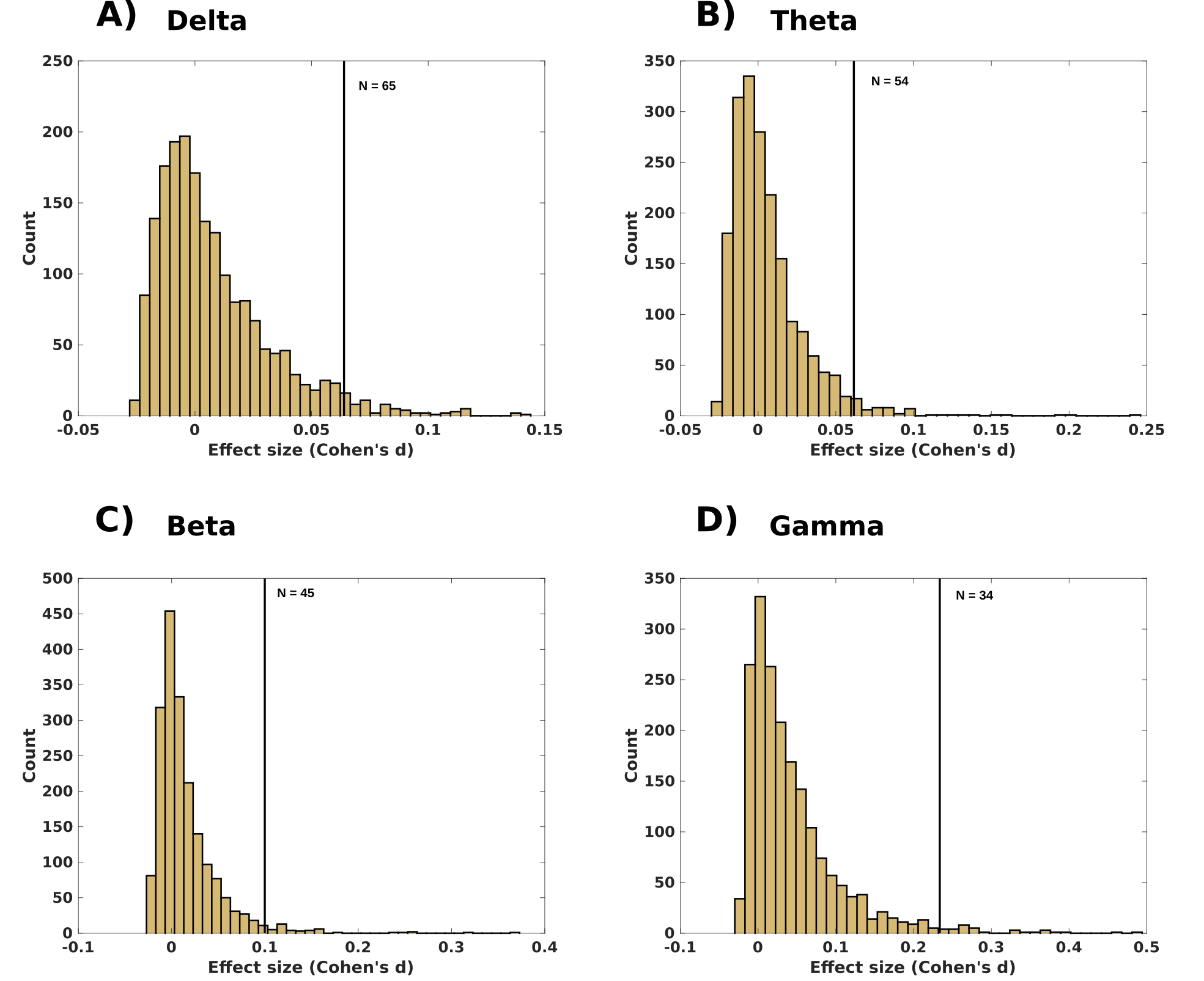

**Fig S7. Histogram of the effect sizes for the four frequency bands.** The vertical line marks the cutoff point for the network of edges showing the largest topic sensitivity. (**A**) Delta band cutoff is at *d* = 0.081 with 65 edges above cutoff. (**B**) Theta band cutoff is at *d* = 0.074 with 54 edges above cutoff. (**C**) Beta band cutoff is at *d* = 0.115 with 45 edges above cutoff. (**D**) Gamma band cutoff is at *d* = 0.234 with 34 edges above cutoff.

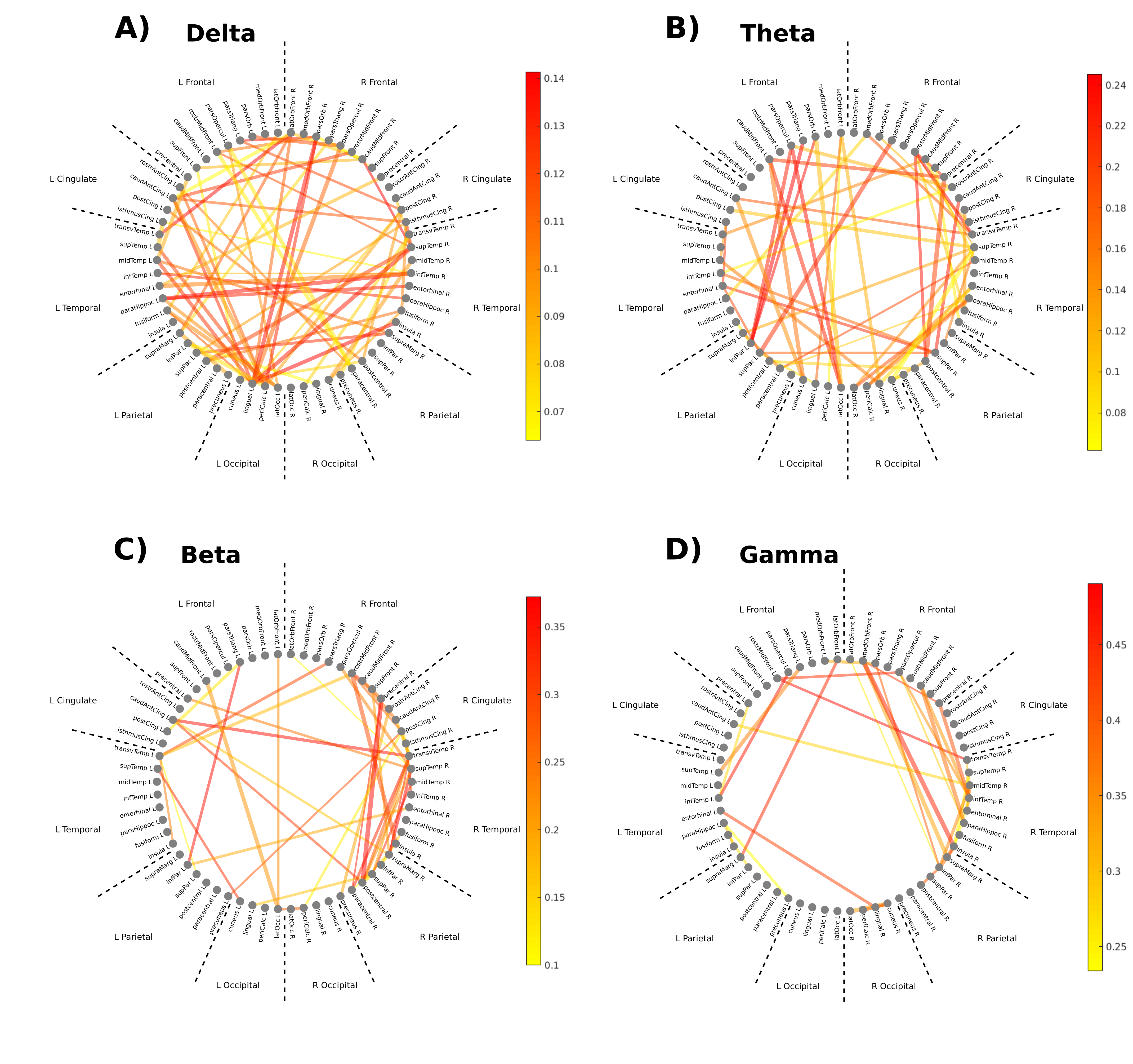

**Fig. S8. Circular diagram of the edges maximizing sensitivity of the similarity difference between within- and across-narratives grouped by anatomical areas, separately for the four frequency bands.** Edge width corresponds to group-level connectivity strength and edge color marks relative contribution from most (red) to lesser (yellow). Note that color mapping (shown right to each panel) differs across the four frequency bands. Panels: (**A**) delta; (**B**) theta; (**C**) beta; (**D**) gamma band.

| **Table S1** |  |  |  |  |
| --- | --- | --- | --- | --- |
| Comparison between within- versus across-topic FC similarity values for the four frequency bands | | | | |
| Frequency band | Within-topic FC matrix similarity | Across-topic FC matrix similarity | Random permutation test results | One-way ANOVA on within-topic similarity values (factor: Topic) |
| **Delta** | *M* = 0.256  (*SD* = 0.041) | *M* = 0.248  (*SD* = 0.041) | est. *p* < .001, *d* = 0.19 | *F*(3, 3116) = 20.58,  *p* < 0.001, *η* = .019 |
| **Theta** | *M* = 0.259  (*SD* = 0.036) | *M* = 0.253  (*SD* = 0.037) | est. *p* < .001, *d* = 0.16 | F(3, 3116) = 23.9,  *p* < 0.001, *η* = .023 |
| **Beta** | *M* = 0.44  (*SD* = 0.032) | *M* = 0.43  (*SD* = 0.029) | est. *p* < .001, *d* = 0.40 | *F*(3, 3116) = 56.36,  *p* < 0.001, *η* = .052 |
| **Gamma** | *M* = 0.39  (*SD* = 0.049) | *M* = 0.35  (*SD* = 0.038) | est. *p* < .001, *d* = 0.94 | *F*(3, 3116) = 85.90,  *p* < 0.001, *η* = .077 |
| *Note.* Effect size *d* refers to Cohen’s *d*. The results of the follow-up comparisons for each one-way ANOVA are depicted in Figure S4. | | | | |

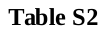

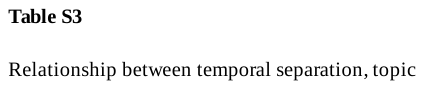

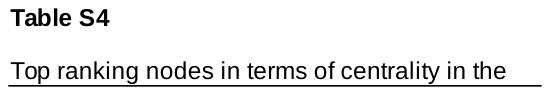

**SI References**

1. O. Szalárdy *et al.*, https://www.biorxiv.org/content/10.1101/2022.02.21.480990v1 (2022).
2. D. H. Brainard, The psychophysics toolbox. Spat. Vis, 10, 433-436 (1997).
3. B. Tóth, F. Honbolygó, O. Szalárdy, G. Orosz, D. Farkas, I. Winkler, The effects of speech processing units on auditory stream segregation and selective attention in a multi-talker (cocktail party) situation. Cortex, 130, 387–400 (2020).
4. B. Tóth, D. Farkas, G. Urbán, O. Szalárdy, G. Orosz, L. Hunyadi, B. Hajdu, A. Kovacs, B. T. Szabó, L. B. Shestopalova, I. Winkler, Attention and speech-processing related functional brain networks activated in a multi-speaker environment. PloS one, 14 (2019).
5. D. J. Barr, R. Levy, C. Scheepers, H. J. Tily, Random effects structure for confirmatory hypothesis testing: Keep it maximal. J. Mem. Lang., 68, 255-278 (2013).
6. D. Bates et al., Parsimonious mixed models. arXiv preprint arXiv:1506.04967 (2015).
7. G. R. Loftus, M.E. Masson, Using confidence intervals in within-subject designs. Psychonom. Bull. Rev., 1, 476-490 (1994).
